# Computational insights into the conformational transition of STING: mechanistic, energetic considerations, and the influence of crucial mutations

**DOI:** 10.1101/2023.05.03.539334

**Authors:** Zhenlu Li, Congran Yue, Shangqiang Xie, Sai Shi, Sheng Ye

## Abstract

STING (stimulator of interferon genes) is a crucial protein in the innate immune system’s response to viral and bacterial infections. In this study, we investigated the mechanistic and energetic mechanism of the conformational transition process of STING activated by cGAMP binding. We found that the STING connector region undergoes an energetically unfavorable transition during this process, which is compensated by the favorable interaction between cGAMP and the STING ligand binding domain. We utilized enhanced sampling methods to study STING’s rotation and finds that several disease-causing mutations, such as N154S and V155L, can result in a smoother transition process, while V147L exhibits unfavorable conformational transition energy. These findings indicate that V147L may not be a gain-of-function mutation, as previously thought, and are further supported by an evolutionary analysis of the STING connector region. Overall, our study provides detailed insights into the mechanism of STING’s rotation and has implications for the development of treatments for STING-related diseases.

## Introduction

The cGAS-STING signaling pathway is an essential component of the innate immune system that plays a critical role in detecting and responding to microbial infections.^1,2^ The pathway is activated by the recognition of cytosolic DNA by the sensor protein cGAS, which leads to the production of cyclic GMP-AMP (cGAMP).^3-6^ cGAMP then binds to and activates the downstream effector protein STING.^7^ Activated STING then recruits and activates the kinase TBK1, which phosphorylates and activates Interferon Regulatory Factor 3 (IRF3).^8,9^ IRF3 translocates to the nucleus and induces the expression of type I interferons and other cytokines, which play a critical role in host defense against viral and bacterial infections.^10,11^

Dysregulation of the STING protein has been associated with multiple autoimmune and inflammatory disorders, highlighting its critical role in maintaining immune homeostasis. Some STING mutations can lead to a loss of function, hindering the body’s ability to mount an effective response against viral infections, while other mutations can result in a gain of function, triggering an overactive immune response and chronic inflammation. SAVI, an exceedingly rare autoinflammatory disease, is a known example of a disease caused by STING mutations at residues 147, 154, and 155. In SAVI, the mutated STING protein is persistently activated, leading to chronic inflammation, blood vessel damage, and organ dysfunction. Given the potential implications of STING in various diseases, it has emerged as a promising therapeutic target for treating illnesses such as cancer, autoimmune disorders, and viral infections.^12-17^

Structural studies have yielded significant insights into the activation mechanism of STING.^18–20^ STING forms a butterfly-shaped dimer, with each subunit consisting of two domains: a ligand-binding domain (LBD, residues 157–379) and a transmembrane domain (TMD, residues 1 to 140). A two-turn connector helix (residues 137–149) and a connector loop (residues 150–156) connect the two domains. The dimeric LBD undergoes a conformational change upon ligand binding of cyclic GMP-AMP (cGAMP), rotating about 180° from its inactive to active state (Fig. 1a). This activated STING subsequently triggers various downstream signaling pathways, including the type I interferon pathway.

**Figure 1:**
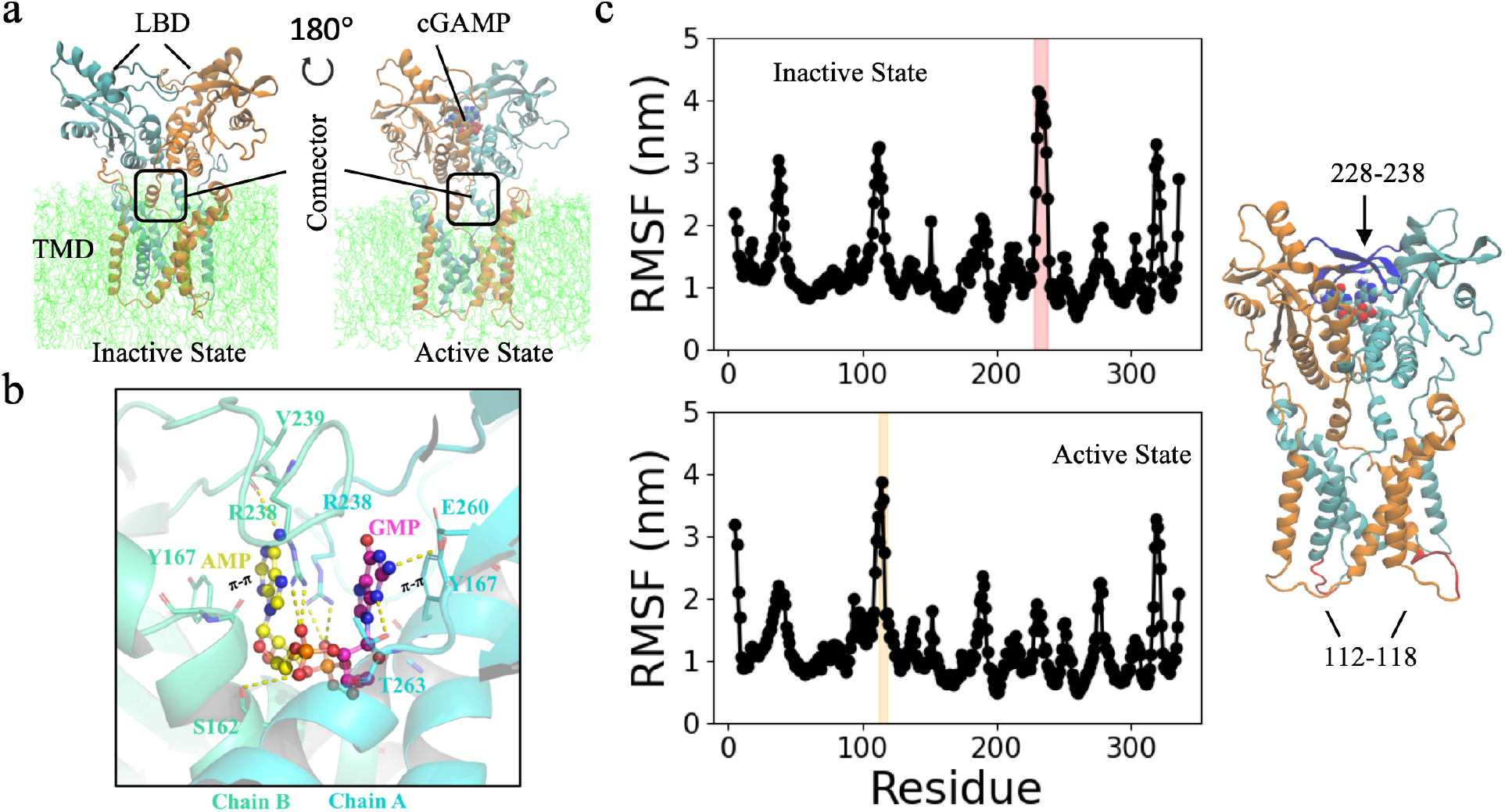
Two kinds of STING conformations. (a) Structural domains of STING at inactive state and active state. (b) cGAMP-STING binding sites. (c) Root mean square fluctuation (RMSF) of STING residues at active and inactive state. Positions of residues 228 to 238 and residues 112 to 118 are highlighted.

Although we have gained a considerable amount of knowledge regarding STING activation, there is still a lack of understanding regarding the specific role of each component in the transition process and the impact of critical mutations. In this study, we utilized atomistic simulations to uncover a more detailed mechanism of STING transition, including the energetic and mechanistic factors driving the process.

## Methods

### Molecular dynamics simulation

The STING protein sequence covers residues 1 to 336, with the C-terminal tail (residues 337 to 379) excluded from simulation due to experimental limitations. The inactive human STING structure was obtained from PDB 6nt5, and the active human STING structure was derived from PDB 6n7t (for chicken) and PDB 4ksy (human STING extracellular domain). cGAMP was built by patching a GMP and AMP molecule in CHARMM force field.^21^ The STING membrane complex was built with CHARMM-GUI web sever.^22^ The membrane is composed of 298 POPC lipids. The protein-membrane is solvated by TIP3P water molecules.^23^ We conducted conventional molecular dynamics simulations to investigate inactive and active states of STING, as well as mutants, for 100-300 ns. Each simulation was repeated for three times. Detailed simulation systems are listed in Table S1. Interactions were calculated using a 1.2 nm cut-off, and 150 mM NaCl ion concentration was used. Simulations were performed at 310 K with the NAMD/2.12 package, and we used the CHARMM36 force field throughout.^24,25^

### Enhanced sampling simulations

We employed targeted molecular dynamics (TMD) simulations to investigate STING rotation. TMD is a powerful approach for inducing conformational changes in known structures by applying geometrical constraints in a time-dependent manner.^26^ To induce STING rotation, we used a force constant of 100 kcal/mol/ in our TMD simulations, and we ran simulations for 20 ns. We applied a tcl rotation force at the beginning of the simulation (1 ns) to guide STING rotation in a clockwise direction from inactive to active state (top view). We also performed TMD simulations in the opposite direction to study STING rotation from active to inactive state, which occurs in an anticlockwise direction.Visualization and analysis of model feature were performed by VMD and Open-source Pymol.

### Simulation analysis

We studied the potential interactions between cGAMP and an inactive STING. To do this, we placed cGAMP onto an inactive STING by superimposing the extracellular domain of STING in its active and inactive states. We then moved cGAMP by 0.7 nm in the Z direction, so that the phosphate group of cGAMP was in the same plane as the guanidine group of arginine. To calculate forces, we extracted R238-cGAMP coordination and calculated the force based on electrostatic potential using a custom Python script. We separated the force into inner and tangential components and neglected van der Waals interactions. We also conducted standard analyses, including pairwise interactions and residue-residue contact maps, using VMD and NAMD script.^27^ In Fig. 4c and Fig. 5b, we binned energy every 0.05 nm of RMSD_1.

### Evolutionary analysis

We conducted an evolutionary analysis by gathering connector region sequences from 20 different species. The amino acids of 135-173 were selected for doing the analysis. To infer a phylogenetic tree, we selected Monosiga brevicollis as an outgroup and performed sequence alignment using mafft.^28^ We used RAxML^29^ to construct the phylogenetic tree and tested 1000 bootstrap tries. In Fig. 5c, we removed the outgroup and aligned the sequences to identify conserved regions. Further details could refer to the study by Starr et al. 2022.^30^

## Results and Discussion

### cGAMP binding provides driven force promoting the rotation of STING

cGAMP is made up of phosphodiester linkages that bind the two nucleotides adenosine monophosphate (AMP) and guanosine monophosphate (GMP). The two adenine nitrogenous bases engage with Y167 of both subunits of STING, whereas the phosphate groups in cGAMP have significant negative potential and interact with R238 of both monomers (Fig. 1b). The atomistic simulations of inactive and active STING (Fig. S1 and Table S1 for simulation systems) indicate that the protein structure is stabilized by the binding of cGAMP, which results in less fluctuation of amino acids in and around the binding area, in particular for the loop from residues 228 to 238. Notably, there is a rise in fluctuation in the intracellular loop of residues 112 to 118, which is not involved in cGAMP binding (Fig. 1c).

Upon binding to cyclic GMP-AMP (cGAMP), the STING protein undergoes a conformational change that both STING monomers rotate inwards towards the ligand-binding pocket. This conformational change brings the main cGAMP-binding amino acids, two R238 and two Y167, closer together while simultaneously pushing apart the two G158 residues to better accommodate cGAMP binding. In the inactive state, the R238-R238 and Y167-Y167 distances of the two STING subunits are 1.57 ± 0.08 nm and 1.44 ± 0.03 nm, respectively, as shown in Fig. 2a. These distances are larger than those in the active state, which are only 1.21 ± 0.36 nm and 1.05 ± 0.06 nm, respectively. In contrast, the G158-G158 distances are increased from 0.44 ± 0.03 nm to 0.93 ± 0.08 nm (Fig. 2a).

**Figure 2:**
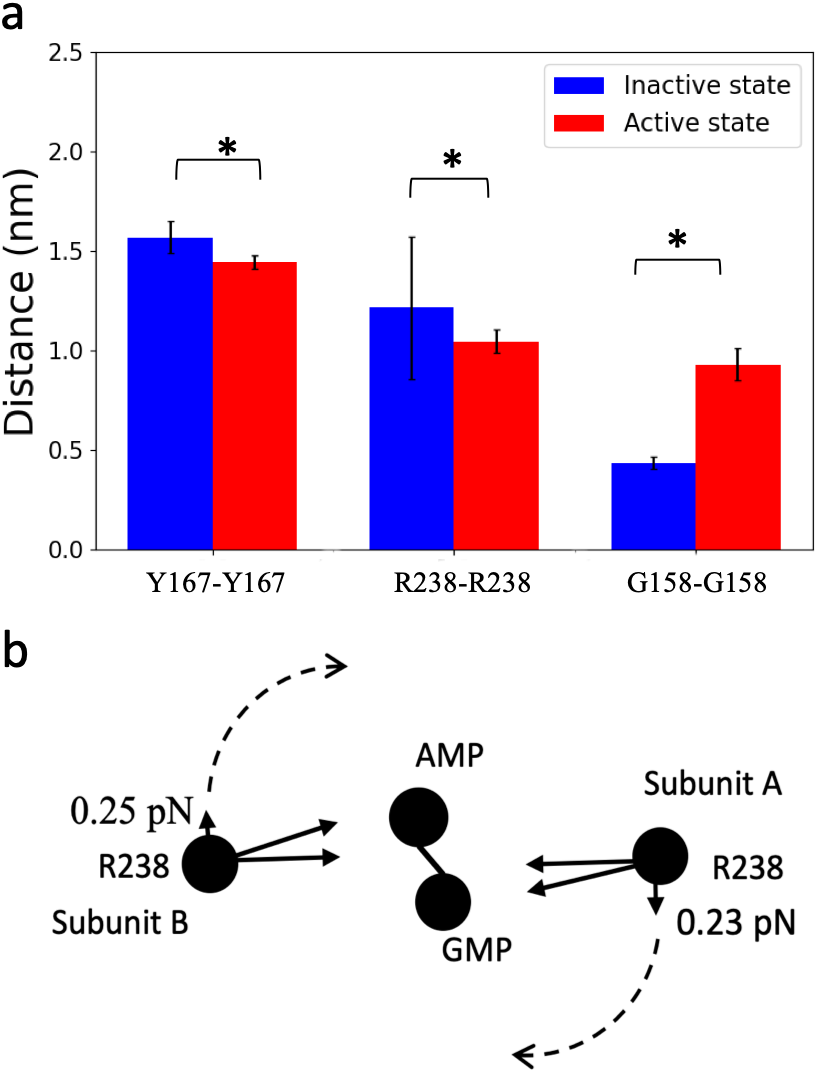
Mechanistic analysis on cGAMP binding and STING rotation. (a) Distance of Y167-Y167, R238-R238, and G158-G158 (*p-value <0.001). (b) Schematic models of tangential force generated by two assymeteric R238-cGAMP interactions. Black filled circle represents center of mass of R238, AMP and GMP.

By superimposing the active and inactive states of the extracellular domain (residues 174-336), it is possible to observe how cGAMP may interact with STING residues in the inactive state. We roughly moved the cGAMP in the Z direction to place it where the phosphate group of cGAMP is in the same plane as the guanidine group of arginine. Specifically, in the inactive state, the phosphate group of each cGAMP component (GMP, AMP) attracts its own R238 partner, resulting in two tangential forces of ∼0.23–0.25 pN that are perpendicular to the R238-R238 connecting line (Fig. 2b). These forces provide the source of power for the rotation of the dimer in the binding pathway.

### The energetics of the STING connector region in two states

In the transition of STING rotation, the most significant conformational change occurs in the the region spanning residues 137 to 173, which can be further divided into the connector helix (137-149), connector loop (150-156), and helix 1 of the LBD domain (157-173). Here, we refer to this region as the connector region. At P173, the helix undergoes a significant kink, which changes the direction of helix 1. There are rare interaction between two monomers for the remaing LBD domain (residues 174 to 336), but the rotation of the connector region would dirve the rotation of entire LBD domain. Comparison of the active and inactive states reveals that in the inactive state, helix 1 of the LBD is interleaved, whereas it becomes two parallel helices in the active state. As a result, inter-helical interactions change significantly. In the inactive state, many inter-helical interactions, particularly between residues N154 and H157 of helix A, and residues E149 and N152 of helix B, form a hydrophilic region. Another important inter-helical interaction involves residues F153 and V155 of helix A and residues L159 and W161 of helix B, generating a hydrophobic region that is outward-facing (Fig. 3a, b).

**Figure 3:**
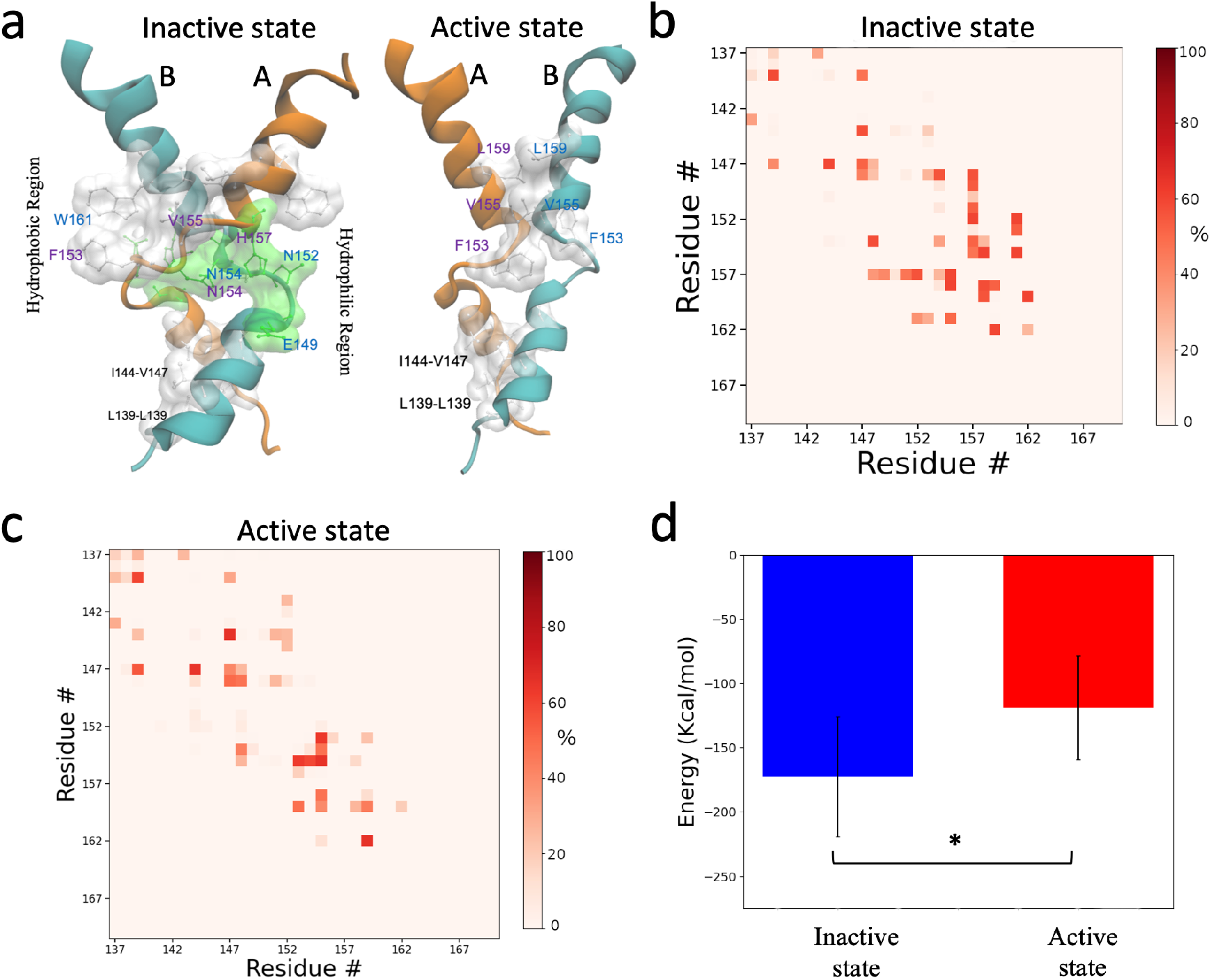
Conformations and energetics of STING connector region. (a) Conformations of STING connector region (residues 137 to 173) in inactive state (left) and active state (right). Highlighted residues represent hydrophobic contact or hydrophilic interactions. Residue-residue contact maps for the (b) inactive state and (c) active state. (d) Comparison of pair interactions for inactive state and active state (*p-value <0.001).

In contrast, in the active state, the inter-helix contact mode is completely different (Fig. 3a, c), resembling a coiled-coil. The former hydrophilic regions are lost, and the former hydrophobic region is largely changed. Inward-facing interactions between pairwise residues, including F153, V155 and L159, maintain hydrophobic interactions. Comparison of the pair interactions of the two states also reveals significant differences. The energy in the active state is -118 ± 40 kcal/mol, much lower than that in the inactive state of -172 ± 46 kcal/mol (Fig. 3d). The inactive-state transition is energetically unfavorable regarding the connector region, but this energy should be largely compensated by the binding of substrate cGAMP molecules. The calculated pair energy of cGAMP-STING is -407 ± 38 kcal/mol.

### Simulations of STING conformation transtion

In order to gain a better understanding of the STING transition process, we conducted targeted molecular dynamics simulations to simulate the transition of STING from an inactive state to an active state, and vice versa (as shown in Fig. 4a). To ensure the correct rotation of STING, we applied an external rotational force at the beginning of the simulation. The evolution of RMSD (root mean square deviation) towards the starting conformation and final conformation is presented in Fig. 4b, which indicates that under the influence of external force, the extracellular domain undergoes a complete 180-degree rotation, resulting in a full transition from one state to the other.

Upon further analysis, we have examined the energy levels at various RMSD positions, shedding light on the transition from the inactive to the active state. The journey from quiescence to activation is not a straightforward ascension of energy. When the configuration deviates from the inactive state by 0.2 nm of RMSD, the energy sharply surges. However, as the configuration progresses, the energy plummets to a low state before surging once again to the energy level representing activation (Fig. 4c). Consequently, we can infer that the active state is an energetically unfavorable state when compared to the inactive state, and the latter is more stable when not in association with cGAMP.

**Figure 4:**
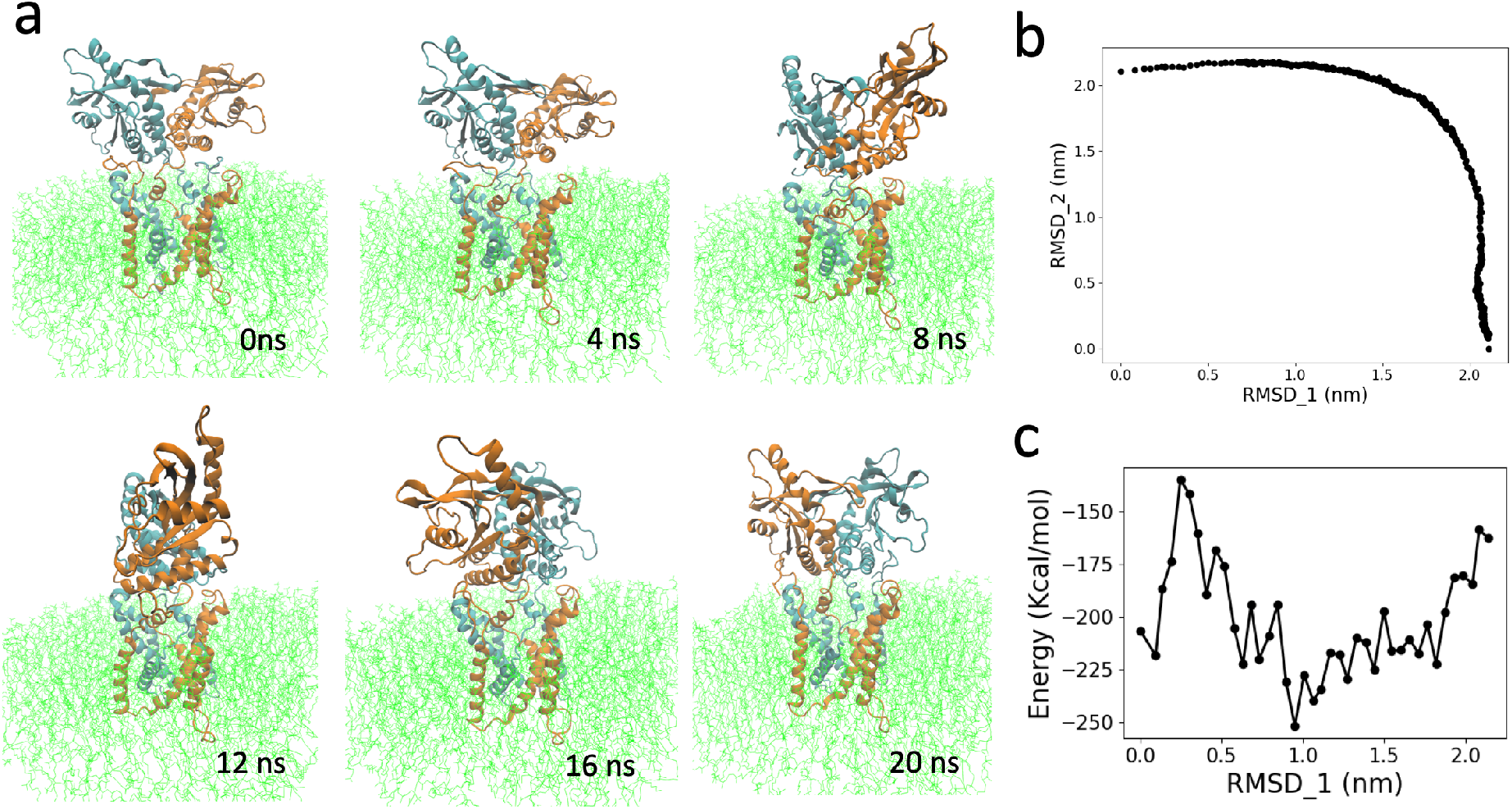
STING rotation revealed by targeted molecular dynamics simulation. (a) Snapshots of STING rotation (here from inactive state to active state). (b) RMSD toward to inactive state (RMSD_1) and active state (RMSD_2) respectively in one representative simulation. (c) Energy of pairwise interaction between residue 137-170 of helix A and B as a function of RMSD_1.

### Influence of mutations on STING rotation

The mutations V147L, N154S, and V155M exert a significant impact on STING activation. To further investigate this effect, we simulated the active and inactive states of STING with these mutations. Our analysis revealed that both N154S and V155M mutations reduce the energy difference between ative state and inactive state in the connector region. The energy of the active and inactive states is -180 ± 50 kcal/mol and -138 ± 36 kcal/mol for N154S mutation, and -193 ± 36 kcal/mol and -167 ± 44 kcal/mol for V155M mutation (Fig. 5a). In the active state, the smaller size of Serine leads to a closer distance between the two helix 1, potentially increasing interactions in the active state. In contrast, the Methionine with its longer branch than Valine, as seen from simulation trajectories, allows it to interact with neighboring F153 and L159, increasing the stability of the active state conformation (Fig. S2).

Previously identified in the mutation maps of SAVI disease patient, V147L was considered a potential gain of function mutation. However, our analysis showed that V147L actually increased the energy of the inactive state (−196 ± 43 kcal/mol), while having less influence on the active state (−117 ± 48 kcal/mol), as a result increasing the energy difference between inactive state and active state (Fig. 5a). Simulation of conformational transitions revealed that the V147L mutation alters the shape of energy curve, but has less impact regarding the absolute energy barrier between inactive and active state (Fig. 5b). In contrast, N154S and especially V155M both display a smoother transition curve and lower energy barrier (Fig. 5b). Therefore, N154S and V155M mutations have the potential to lead to higher levels of self-activation of STING, even in the absence of ligand binding. However, the V147L mutation may not have this effect.

**Figure 5:**
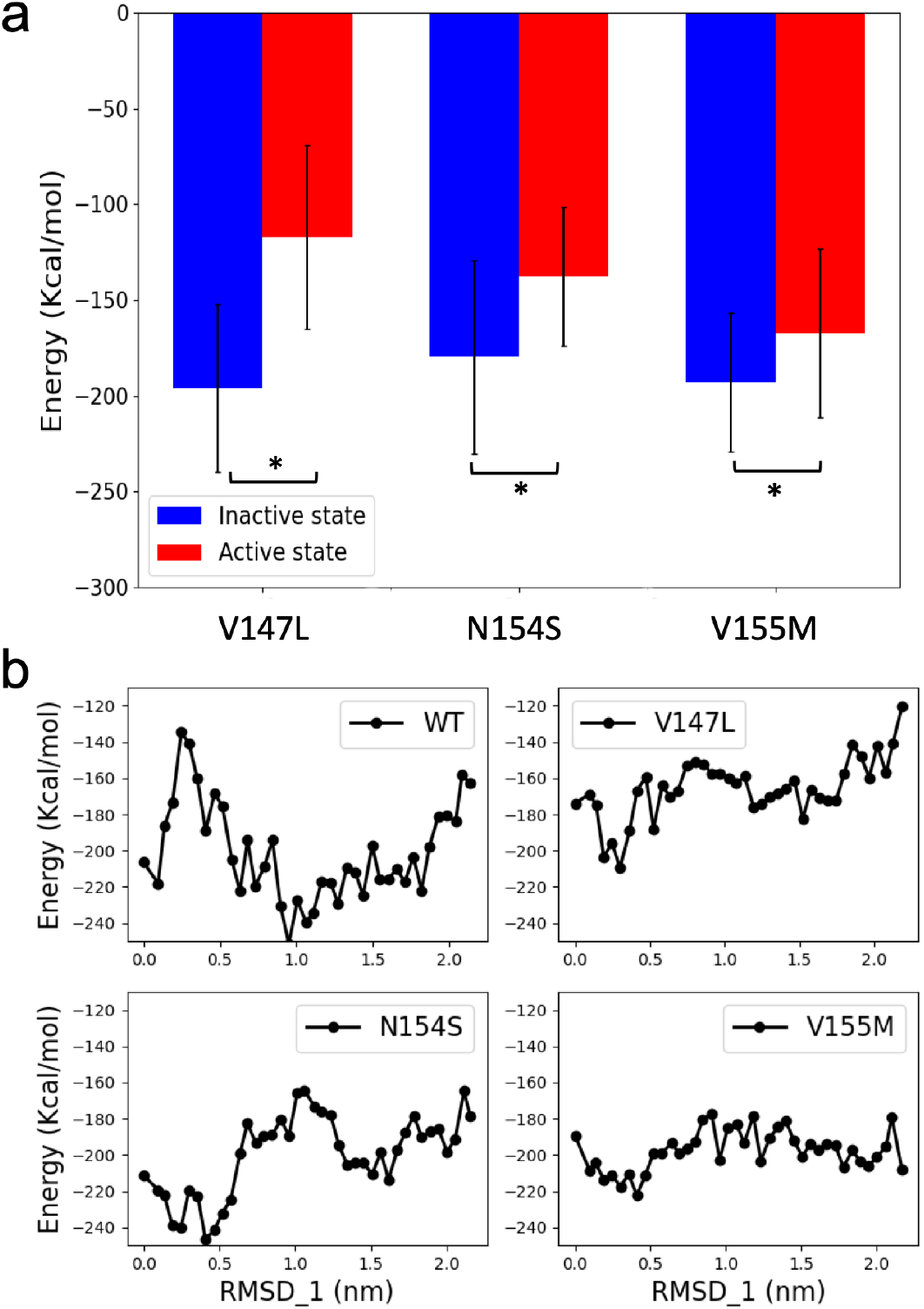
Influence of mutations on STING rotations. (a) Comparison of pair interactions of the connector region for inactive state and active state for three STING mutants (*p-value <0.001). (b) Energy-RMSD_1 plots during transition process from inactive state to active state for wild type STING and three mutants.

### Evolutionary insights into the connector region

The cGAS-STING pathway has deep evolutionary roots, with the STING ligand binding domain (LBD) and its unique cyclic dinucleotide binding function remaining conserved for over 600 million years.^31,32^ However, the appearance of the transmembrane region (TMD) is a feature unique to animal phyla, including unicellular choanoflagellates, some invertebrate insects, fish, birds, and mammals. In bacteria and certain animals, such as in Lophotrochozoans, the TMD is replaced by a TIR domain (Fig. S3). To investigate the evolutionary pattern of the connector regions, we focused on the STING connector region of representative species of animal phyla, conducting sequence alignment and a phylogenetic tree analysis (Fig. 6a,b).

**Figure 6:**
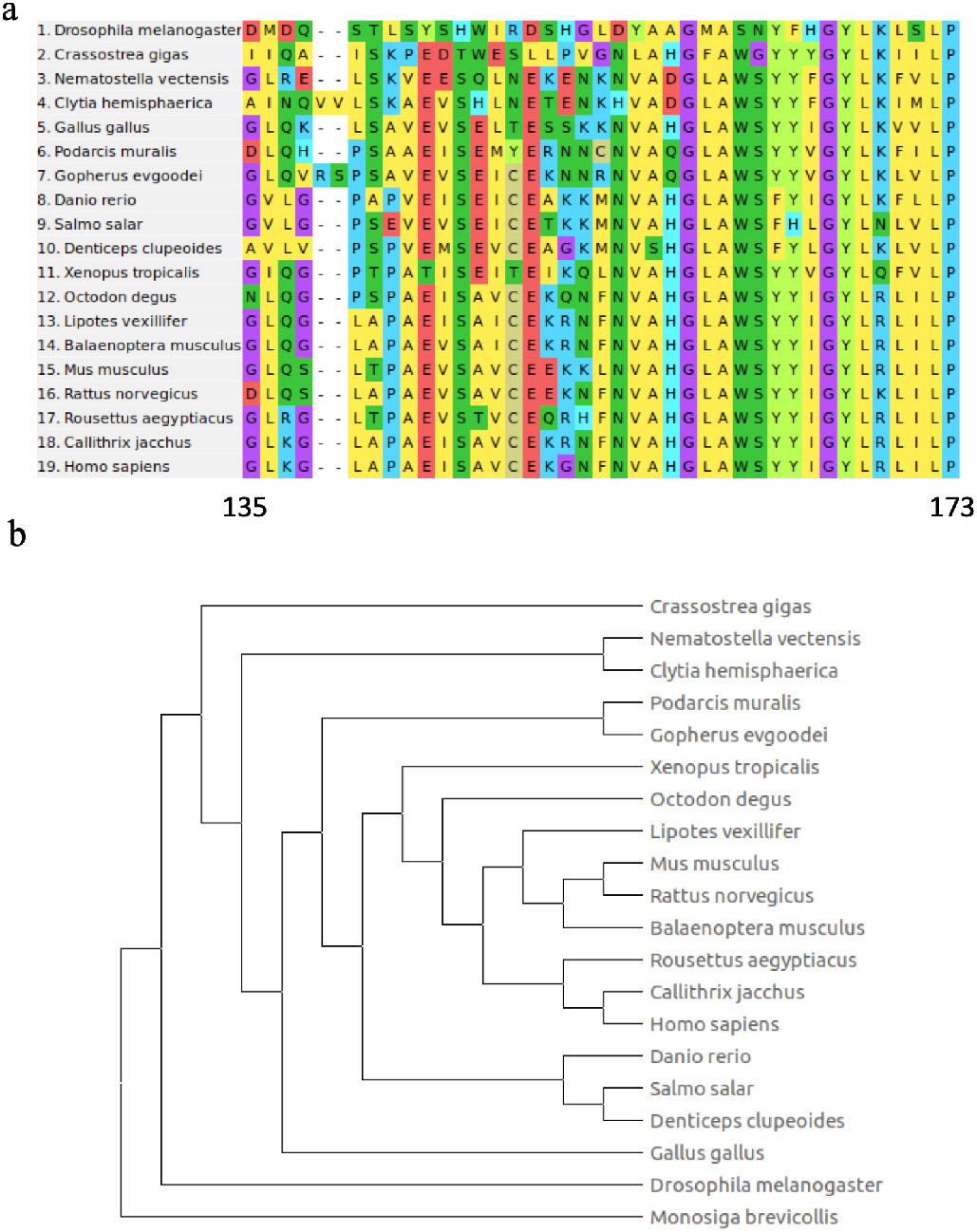
Evolutionary analysis of the STING connector region. (a) Alignment of the connector regions from 19 animal phyla, illustrating sequence conservation and variation. (b) Phylogenetic tree constructed using the connector region sequences from the 19 animal phyla, with the sequence of monosiga brevicollis serving as an outgroup to root the tree.

Analysis of diverse proteins from invertebrates to humans showed that the kink region (proline 173) is conserved, along with several other residues such as G158, A160, G166, Y167, L168 and L172 (Fig. 6a). These amino acids either interact with cGAMP (Fig. 1a) or are adjacent to cGAMP, making them likely to be conserved. In contrast, the pre-connector region and the beginning of the connector region (residues 135 to 142) exhibited significant amino acid diversity and were not highly conserved. Fruit flies and several kinds of marine organisms, such as Crassostrea gigas, Nematostella vectensis, and Clytia hemisphaerica, showed the most distinct sequence differences compared to the sequence of Homo sapiens (Fig. 6b). Interestingly, the sequence of chicken diverges quite early in the phylogenetic tree, even before fishes and amphibious animals, potentially indicating some special feature of STING in birds.

The middle of the connector region, consisting of amino acids 143 to 157, is critical for STING rotation, as discussed above. Notably, N154 and V155 were highly conserved, whereas V147 was not fully conserved. V147I or V147L mutations are frequently observed in multiple species, including higher animals such as chicken and mouse. If these mutations confer a gain-of-function, they may lead to uncontrolled self-activation of the STING system, which could be harmful to these species. While some literature suggests that V147 is associated with gain-of-function disease and potentially related to STING self-activation, our evolutionary and energy analyses indicate that V147 mutation may not significantly reduce the STING transition energy. Therefore, we speculate that V147 mutation is unlikely to be related to STING self-activation.

## Conclusions

Insights into the conformational transition mechanism of STING molecules have been provided by this study. The connector region of STING exists in two forms: an energetically favorable inactive form and a less favorable active form. However, the binding of the ligand cGAMP compensates for the loss of energy. Mechanistically, when cGAMP binds to STING, the interactions between cGAMP and two arginine generate a tangential force that rotates the ligand binding domain of STING. At the same time, the STING rotation creates more space for optimal cGAMP binding. Notably, two disease-causing mutations, N154S and V155M, reduce the transition energy, promoting the transition from the inactive to active state. However, V147L has the opposite effect on the energy transition. Evolutionary analysis of the connector region shows that V147 is not highly conserved amino acid, which indicates that V147L may not be a gain-of-function mutation as previously thought. This knowledge can aid in understanding the activation mechanism of STING and guide the design of drugs targeting STING.

## Acknowledgements

This work was supported in party by Ministry of Science and Technology (2020YFA0908500 to S.Y.) and the National Natural Science Foundation of China (31971127 to S.Y). L.Z.L was supported by start-up funding.

## Author Contribution

**L.Z.L, Y.S:** Conceptualization, **L.Z.L, X.S.Q**., **Y.C.R. S.S**: Data curation, Investigation, Writing-Original draft preparation. **L.Z.L**: Modeling, Visualization. **L.Z.L**., **Y.S**.: Supervision., **L.Z. L**., **Y.S:** Writing-Reviewing and Editing.

## Data Availability

All data are available in the paper and the Supplementary Data.

