## Supplementary_Information for "Computational insights into the conformational transition of STING: mechanistic, energetic considerations, and the influence of crucial mutations"

Table S1: List of simulation systems.

| Conventional Molecular Dynamics Simulation | | | |
|---|---|---|---|
| | | Time (ns) | Repeats |
| STING WT | | 250 ns | 3 |
| STING V147L | | 100 ns | 3 |
| STING N154S | | 100 ns | 3 |
| STING V155M | | 100 ns | 3 |
| Enhanced Sampling Simulation | | | |
| STING WT | inactive→active | 20 ns | 3 |
| | active→inactive | | |
| STING V147L | inactive→active | 20 ns | 3 |
| | active→inactive | | |
| STING N154S | inactive→active | 20 ns | 3 |
| | active→inactive | | |
| STING V155M | inactive→active | 20 ns | 3 |
| | active→inactive | | |

Figure S1: Time evolution of RMSD for 3 independent simulations for (a) inactive state and (b) active state.

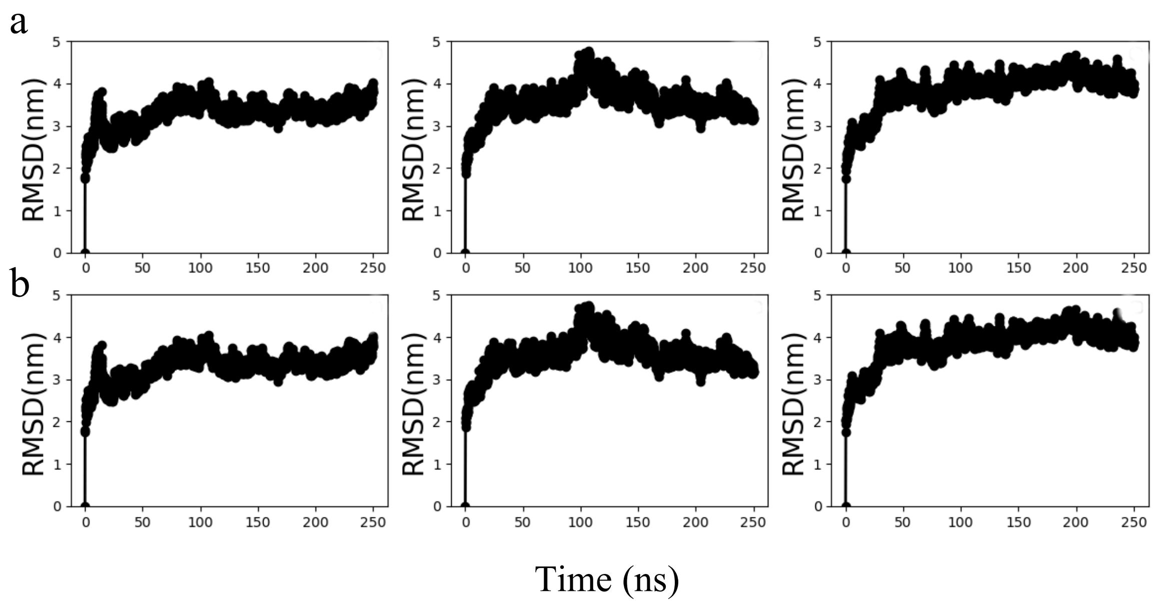


Figure S2: The interactions of M155 with neighboring F153 and L159 in the active state.

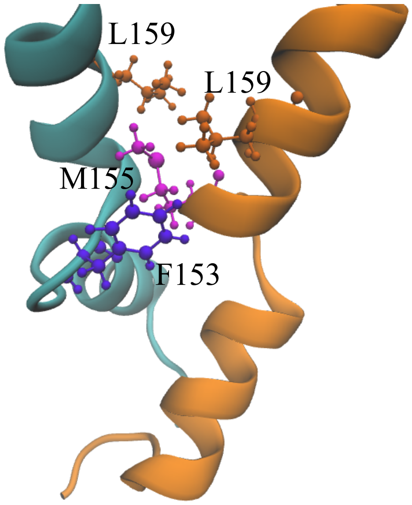

Figure S3: Evolution of STING structural domains.

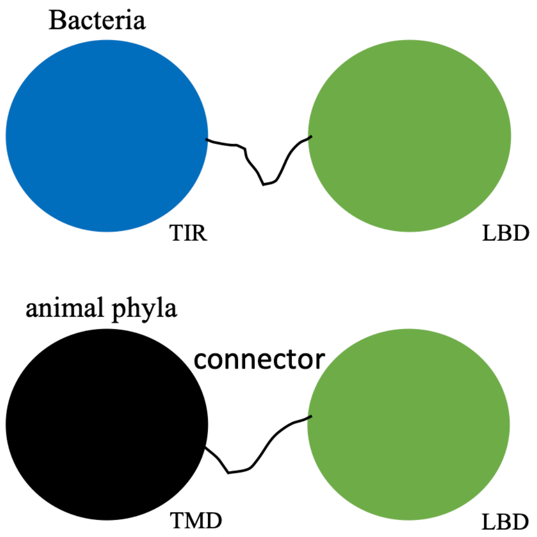
